# Distinct roles of the major binding residues in the cation-binding pocket of MelB

**DOI:** 10.1101/2024.02.27.582382

**Authors:** Parameswaran Hariharan, Amirhossein Bakhtiiari, Ruibin Liang, Lan Guan

**Affiliations:** Department of Cell Physiology and Molecular Biophysics, Center for Membrane Protein Research, School of Medicine, Texas Tech University Health Sciences Center, Lubbock, TX; Department of Chemistry and Biochemistry, Texas Tech University, Lubbock, TX

**Keywords:** symport, bioenergetics, PEF, ITC, protonation, cation selectivity, external pH, melibiose permease

## Abstract

*Salmonella enterica* serovar Typhimurium melibiose permease (MelB_St_) is a prototype of the major facilitator superfamily (MFS) transporters, which play important roles in human health and diseases. MelB_St_ catalyzed the symport of galactosides with either H^+^, Li^+^, or Na^+^, but prefers the coupling with Na^+^. Previously, we determined the structures of the inward- and outward-facing conformation of MelB_St_, as well as the molecular recognition for galactoside and Na^+^. However, the molecular mechanisms for H^+^- and Na^+^-coupled symport still remain poorly understood. We have solved two x-ray crystal structures of MelB_St_ cation-binding site mutants D59C at an unliganded apo-state and D55C at a ligand-bound state, and both structures display the outward-facing conformations virtually identical as published previously. We determined the energetic contributions of three major Na^+^-binding residues in cation selectivity for Na^+^ and H^+^ by the free energy simulations. The D55C mutant converted MelB_St_ to a solely H^+^-coupled symporter, and together with the free-energy perturbation calculation, Asp59 is affirmed to be the sole protonation site of MelB_St_. Unexpectedly, the H^+^-coupled melibiose transport with poor activities at higher ΔpH and better activities at reversal ΔpH was observed, supporting that the membrane potential is the primary driving force for the H^+^-coupled symport mediated by MelB_St_. This integrated study of crystal structure, bioenergetics, and free energy simulations, demonstrated the distinct roles of the major binding residues in the cation-binding pocket.

## Introduction

The secondary active transporters, including symporters or antiporters, play important roles in physiology and pathology. The major facilitator superfamily (MFS) transporters contain a large group of cation-coupled symporters, most of which couple to either H^+^ or Na^+^ electrochemical gradient. The melibiose permease of *Salmonella enterica* serovar Typhimurium (MelB_St_), a member of MFS transporters, catalyzes the symport of a galactopyranoside with either of H^+^, Li^+^, or Na^+^, MelB is a unique model system for studying cation-coupled transport mechanisms (1-11). The Na^+^-coupled lipids transporter (MFSD2A) expressed in the major barriers, such as the blood-brain barrier or blood-retina barrier, plays a critical role in the uptake of essential lipids into these neural tissues. These transporters shared the same cation-binding site as the bacterial MelB but were much less characterized(9, 10, 12), and MelB as a well-studied model system for this group of transporters is useful for expanding our knowledgebase of cation-coupled transport mechanisms.

Two crystal structures with a bound sugar analog have been reported for a cation-binding site mutant D59C MelB_St_ in its outward-facing conformation, which provided essential information on the determinants for the sugar substrate specificity (8) (**Fig 1a**). Recently, a Na^+^-bound inward-facing conformation of the WT MelB_St_ has been determined by cryoEM single-particle analysis, which reveals the Na^+^ specificity determinants(11) **(Fig 1b)**. Binding affinity measurements of melibiose and Na^+^ or Li^+^ to MelB_St_ in the absence or presence of the other have been extensively carried out via isothermal titration calorimetry (ITC), which revealed positive cooperativity between the sugar and coupling cation with MelB_St_ (13). The cooperativity number, defined as the ratio between the changes in the *K*_d_ values of melibiose and cation in the absence or presence of the other, reflects the coupling efficiency for transporting melibiose with its coupling action. For Na^+^ or Li^+^, the cooperativity number is approximately 8 (13) and 5 (6), respectively, which are significantly greater than that between melibiose and H^+^. It is likely that this number is less than 2 based on the effect of sugar on the H^+^ affinity(13). The structures with bound sugar or Na^+^ reveal that the two binding pockets are in close proximity with no direct overlap (13). The molecular basis for the coupling between the sugar and cation is still elusive.

**Fig. 1.**
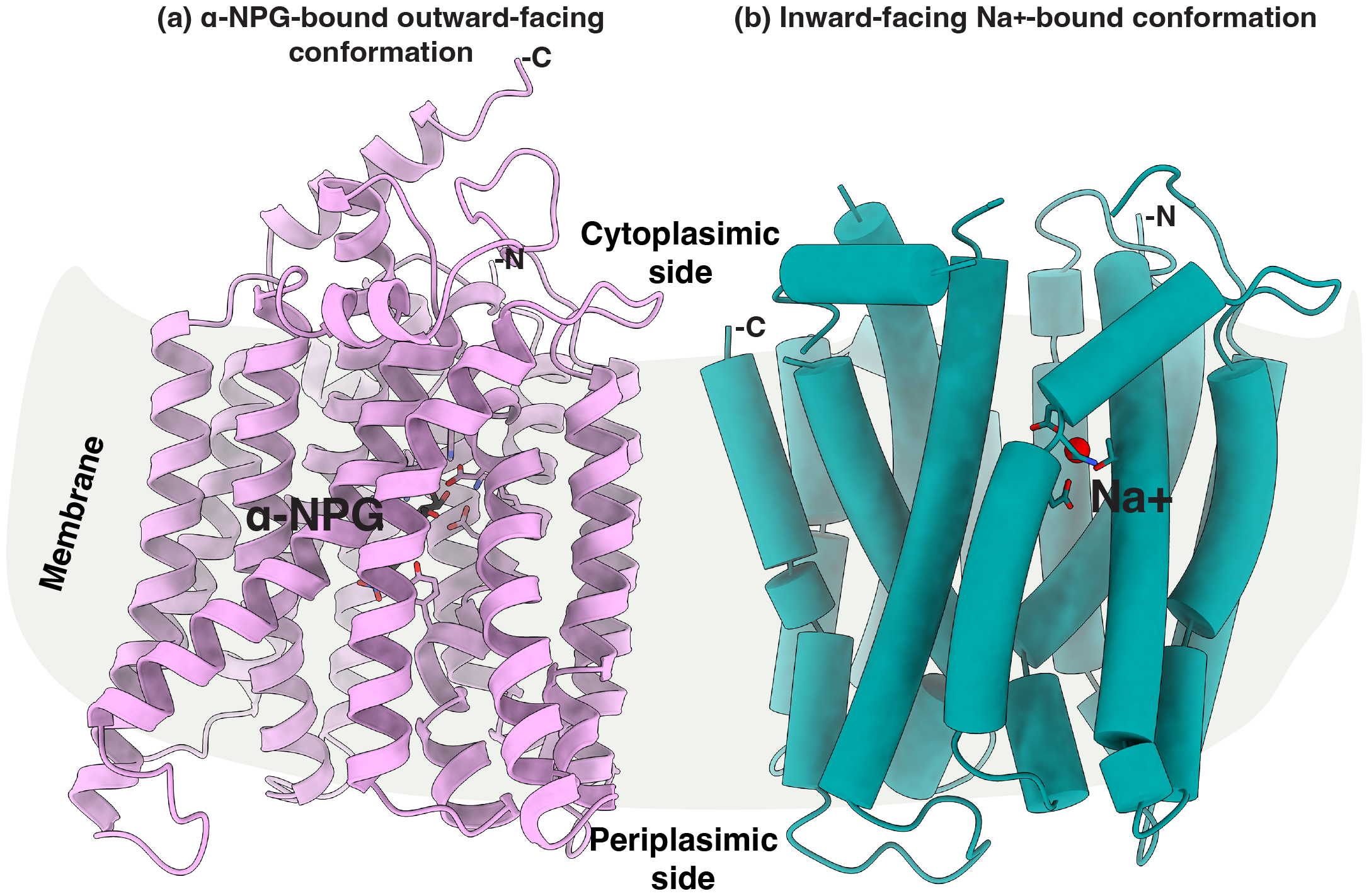
Structures of MelB_St_. **(a)** The ligand-bound outward-facing conformation [PDB ID 7L17]. The cartoon helical representation of the x-ray crystal structure of D59C MelB_St_ mutant with bound α-NPG (8). The sugar-binding sidechains are highlighted in the sticks and α-NPG molecule is colored in black. **(b)** The inward-facing structure of MelB_St_ [PDB ID 8T60](11). Cylindrical helices representation of the cryoEM structure of the WT MelB_St_ bound with Na^+^. The Na^+^-binding sidechains are highlighted in the sticks and Na^+^ is colored in red. The middle loop and the C-terminal tail are disordered. The sidedness of the membrane is indicated. For both structures, the helices II and XI were placed at the front and N-terminus was placed at the right side.

It is noteworthy that the sugar-binding affinity is conformation-dependent(11), in addition to being sensitive to the binding of the coupling cation and its identity. It is usually quite technically challenging to determine the substate-binding affinity for a specific conformation of a transporter because of the conformational variability. The conformation dependency of the sugar-binding affinity of MelB_St_ was obtained by using an inward-facing conformation-selective nanobody Nb725. By trapping MelB_St_ at an inward-facing state, both experimental sugar-binding assay and cryoEM structure support a low-affinity state of the sugar-binding pocket (11, 14).

Remarkably, the Na^+^ binding to the inward-facing conformation remains unchanged (11, 14). All-atom molecular dynamics simulations revealed the virtually identical configurations of this cation binding pocket at either inward- or outward-facing conformation(11). These results undoubtedly verified the stepped binding kinetic model for melibiose/Na^+^ symport at a sequential order of Na^+^ binding first and its release following the sugar release at the other surface, which was constructed based on transport assays, such as efflux and exchange, in a combination of mutants and sugar-binding assay (4, 7, 15, 16).

Both structural and functional studies support that the stoichiometry of the substrate, cation, and transporter is unity. This single cation site for Na^+^ also accommodates H^+^ or Li^+^ since the Na^+^ binding to MelB_St_ is inhibited by Li^+^ (4) or H^+^(13). The stoichiometry number of the coupling H^+^ was verified by measuring the buffer protonation upon the binding of Na^+^ in a set of buffer systems at varying pH values. Structurally, the cation-binding site is formed by two helices within only the N-terminal helical bundle or N-lobe, which might be the structural basis for the unchanged affinity between the outward- and inward-facing states. The Na^+^ binding involves two negatively charged residues, Asp55 and Asp59 (helix II), and two polar residues, Asn58 (helix II) and Thr121 (helix IV). In this shared cation-binding pocket, only Asp55 or Asp59 could serve as a protonation site. The absolute dissolution constant for H^+^, i.e., the p*K*a value, has been determined to be in a slightly acidic range of 6.25 and 6.5 in the absence or presence of melibiose (13).

Asp59 has been suggested to be the protonation site in MelB_St_ for the H^+^-coupled transport (7, 8) and MelB_Ec_ (17, 18). The extensive functional analysis and structure determination show that the D59C mutant loses the active transport of melibiose coupled to the translocation of either Na^+^, Li^+^, or H^+^, but retains the capability of melibiose transport driven by the melibiose concentration gradient(7, 8, 19). D59C mutant loses the binding to Na^+^ or Li^+^(13), and it is a uniporter mutant of MelB_St_.

D55C MelB_St_ also loses the binding affinity with Na^+^ or Li^+^ and is unable to mediate Na^+^- or Li^+^-coupled melibiose active transport. Different from the D59C mutant, the D55C mutant retains the H^+^-coupled melibiose active transport activity(7), which excluded Adp55’s role in the H^+^ binding. The single-site mutagenesis at the Thr121 showed that Ala or Pro replacement only selectively eliminates the Na^+^ binding and Na^+^-coupled melibiose transport but retains the full transport activity coupled to Li^+^ or H^+^(14).

All obtained data suggested the Asp59 as the sole protonation site; however, the molecular basis for the H^+^-coupled transport is unclear. The D55C mutation offered an excellent tool to isolate the H^+^-coupling transport mode to gain insightful knowledge into the H^+^-coupled transport mediated by a Na^+^-coupled transporter without potentially contaminated Na^+^ interference. Here, we report two crystal structures, an Apo D59C mutant and a ligand-bound D55C mutant, and analyze their structures in detail. Furthermore, we performed free energy calculations using molecular dynamics (MD) to quantify the energetic contributions of each of the three side chains (Asp55, Asp59, and Thr121) to the binding of Na^+^ or H^+^, respectively. All data further support the conclusion that Asp59 is the protonation site of MelB_St_ and Asp55 and Asp59 are the major contributors to the Na^+^ binding affinity. Our study of the pH effects on sugar-binding and transport with MelB_St_ showed that membrane potential is the primary driving force for the H^+^-coupled update.

## Results

### Melibiose transport activity

The results of melibiose active transport mediated by the WT MelB_St_ and the mutants D55C and D59C have been published previously (7, 8, 19). The data are reproducible and re-plotted for comparison with the melibiose fermentation data. At a melibiose concentration of 0.4 mM, the WT MelB_St_ expressed in *E. coli* DW2 cells (*mel*A^+^*B*^−^, *lacZ*^−^*Y*^−^) accumulated melibiose in the presence of Na^+^ or Li^+^ at least 6-fold higher than that in the absence of Na^+^ or Li^+^ **(Fig. 2a, red or blue)**. The melibiose transport in the absence of Na^+^ and Li^+^ has been annotated to H^+^-coupled transport (**Fig. 2a, black**). On the MacConkey agar plate containing melibiose as the sole carbon source and neutral red dye as pH indicator for the acidification yielded by melibiose fermentation, the *E. coli* DW2 cells with no MelB grew yellow colonies indicating no melibiose fermentation (**Fig. 2a, inset**) and the presence of WT MelB_St_ changed the color of colonies and agar to magenta indicating good melibiose-downhill transport and fermentation (**Fig. 2a, inset**). The D55C mutant selectively lost the symport activities with Na^+^ or Li^+^, but fully retained the H^+^-coupled melibiose symport activity (**Fig. 2a**). The initial rates of transport and the steady-state accumulation between the WT and the D55C mutant are indistinguishable. The D55C MelB_St_ mutant also reproducibly fermented the melibiose as the WT did (**Fig. 2a, inset**). The uniporter D59C mutant lost all symport activity but retained the melibiose fermentation (**Fig. 2a**) (8).

**Fig. 2.**
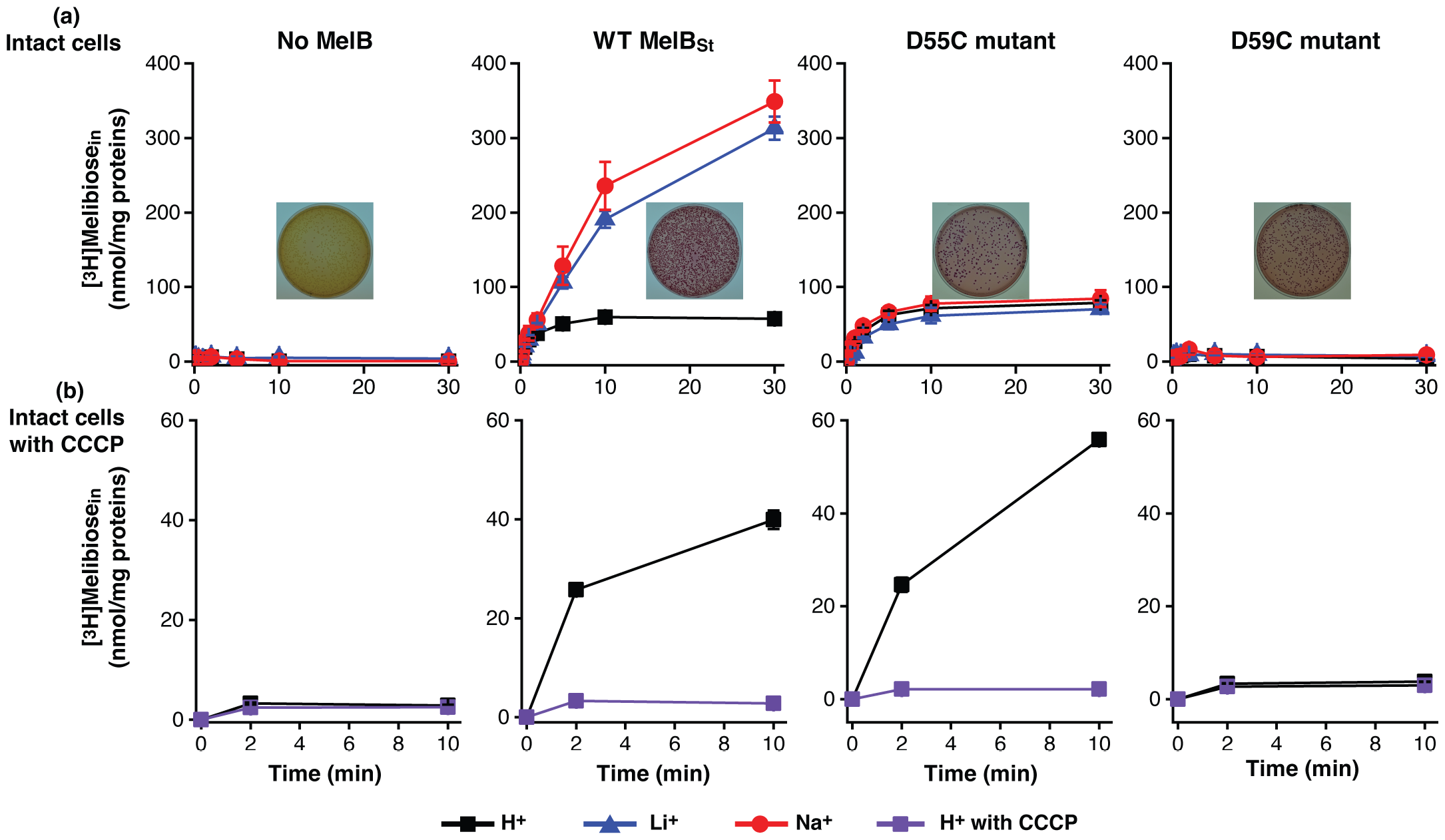
Intact cell transport assay. *E. coli* DW2 cells (*melA*^+^*B*^−^, *lacZ*^−^*Y*^−^) expressing the plasmid-encoding WT MelB_St_ mutants D55C or D59C were prepared for [^3^H]melibiose transport assay. **(a)**Melibiose transport coupled to H, Na. or Li. Melibiose transport was conducted at 0.4 mM (specific activity of 10 mCi/mmol) in the presence of 20 mM Na^+^ or Li^+^ or the absence of Na^+^ and Li^+^ as described in Methods. The cells transformed without MelB were the negative control. **Inset**, melibiose fermentation assay. The cells were also plated onto the MacConkey agar plate containing 30 mM melibiose as the sole carbon source and neutral red dye as pH indicator. The plates were photographed after 16-18 h incubation at 37 °C. Yellow colonies, no melibiose fermentation; magenta colonies, good melibiose fermentation. **(b)** CCCP effect. The H^+^-coupled melibiose uptake at a time points of zero, 2, or 10 m were measured with the cells pre-incubated with 10 μM CCCP. Each test was repeated in 2-3 times.

To further test the melibiose uptake coupled to the electrochemical H^+^ gradient, the melibiose transport assay was conducted with the cells pre-incubated with the proton ionophore, the uncoupler carbonyl cyanide m□chlorophenylhydrazone (CCCP) at 10 μM (**Fig. 2b**). Clearly, CCCP completely inhibited all transport activity at all pH values, confirming the H^+^-coupled melibiose transport mediated by the D55C MelB_St_ and the WT.

### Binding affinity determination by ITC

The D55C mutant has been shown to lose the Na^+^ binding (7, 13). With ITC, Na^+^ or Li^+^ titration at higher concentrations in the absence or presence of saturating concentration of melibiose further confirmed that the D55C mutation lost the binding of Na^+^ and Li^+^ (**Fig. 3a-b**).

**Fig. 3.**
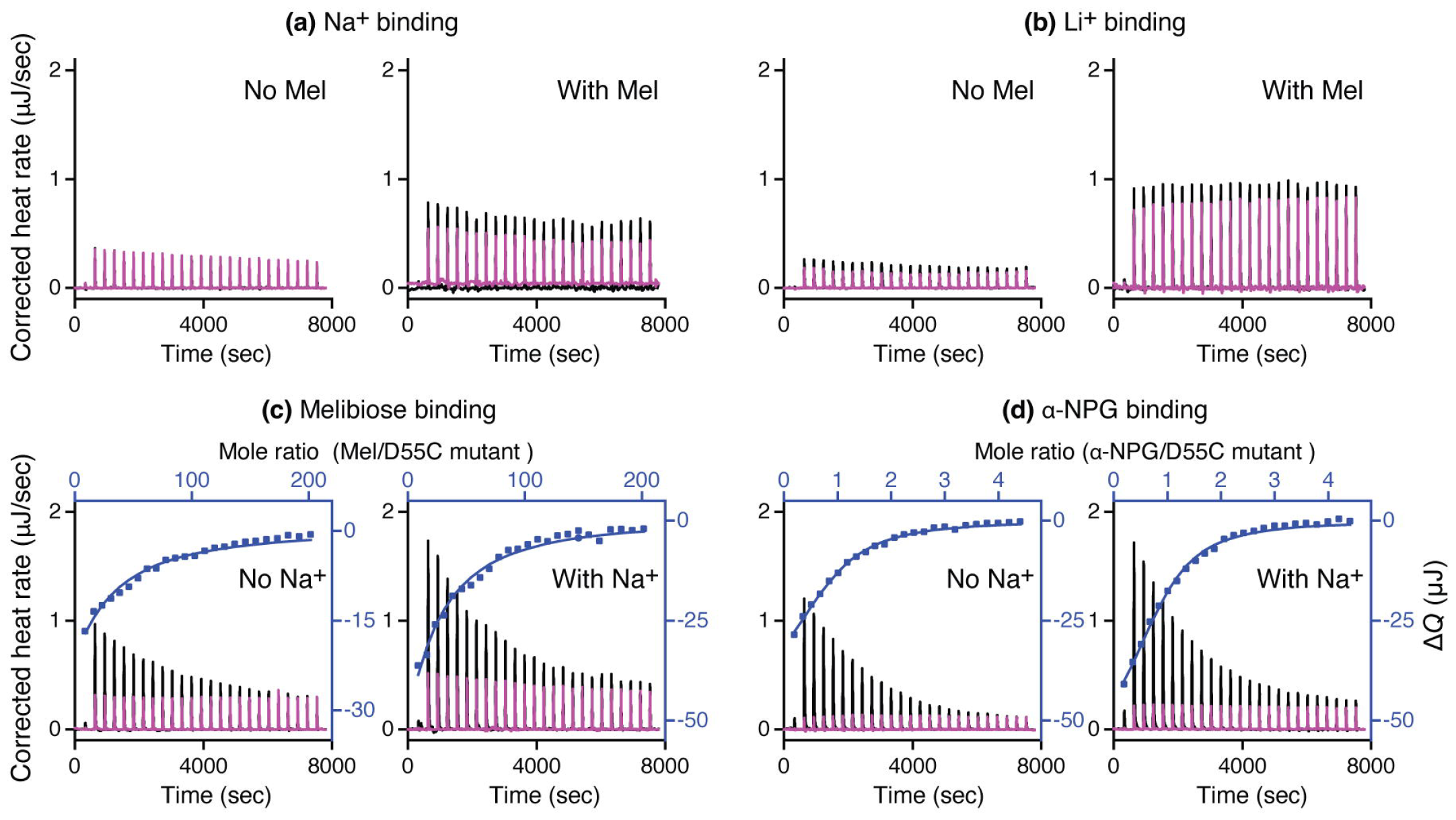
Binding assay via ITC. All binding assays to D55C MelB_St_ mutant were conducted with ITC calorimeters (TA Instruments) at 25 °C. **(a)** Na^+^ binding in the absence or presence of melibiose. The 40 mM NaCl concentration was used for titration, which is 8-fold greater, or 20-fold greater than that used for the WT in the absence or presence of the melibiose, respectively.**(b)** Li^+^ binding in the absence or presence of melibiose. The 40 mM LiCl concentration was used for titration, which is 8-fold greater, or 20-fold greater than that used for the WT in the absence or presence of the melibiose, respectively. **(c)** Melibiose binding in the absence or presence of Na^+^. **(d)** α-NPG binding in the absence or presence of Na^+^. The thermogram was plotted as the baseline-corrected heat rate (μJ/sec; left axis) vs. time (bottom axis) for the titrant to MelB_St_ (black) or to buffer (red) under an identical scale. The heat change Δ*Q* (μJ; filled blue symbol) was plotted against the ligand/D55C mutant MelB_St_ molar ratio (top/right axes).

A previous qualitative Trp to dansyl-galactoside (D^2^G) FRET assay suggested that the D55C mutation binds melibiose and D^2^G(7). The quantitative ITC measurements showed that melibiose binding to D55C mutant in the absence of Na^+^ or Li^+^ exhibits a *K*_d_ value of 5.82 ± 0.46 mM (**Table 1; Fig. 3c**), which is significantly lower than the published value of 9.28 ± 0.23 mM for the WT (8) under the same condition (P<0.01). In the presence of Na^+^ or Li^+^, the WT exhibited an 8.51-fold decrease in the *K*_d_ value for melibiose, but no change was observed with the D55C mutant, which is consistent with no binding of Na^+^ or Li^+^(7).

**Table 1.** *K*_d_ of sugar binding to MelB_St_.

| | pH | Na<sup>+</sup><br>(mM) | Melibiose (mM) | Affinity<br>change | α-NPG<br>(μM) | Affinity<br>change |
| --- | --- | --- | --- | --- | --- | --- |
| WT<sup>^</sup> | 7.5 | / | 9.28 ± 0.23<sup>*#</sup><br>(n = 3) | +8.51-fold | 43.20 ± 0.62<br>(n = 2) | + 1.68-fold |
| | 7.5 | 100 | 1.09 ± 0.06<br>(n = 4) | | 25.69 ± 4.42<br>(n = 2) | |
| D55C | 7.5 | / | 5.82 ± 0.46<sup>#</sup><br>(n = 2) | +1.14-fold | 15.06 ± 1.86<sup>#</sup><br>(n = 2) | +1.11 |
| | 7.5 | 100 | 5.09 ± 0.15<br>(n = 2) | | 13.54 ± 1.38<br>(n = 2) | |
| WT | 6.25 | / | 6.19 ± 0.59<sup>\$</sup><br>(n = 2) | +1.37-fold | | |
| WT | 8.5 | / | 8.50 ± 0.21<sup>\$</sup><br>(n = 3) | | | |

<sup>\*</sup>sem, standard error; n, the number of tests.
<sup>^</sup>, data published in reference of (8).
<sup>#</sup>, unpaired t-test, P<0.01.
<sup>\$</sup>, unpaired t-test, P<0.05.

**Table 2.**
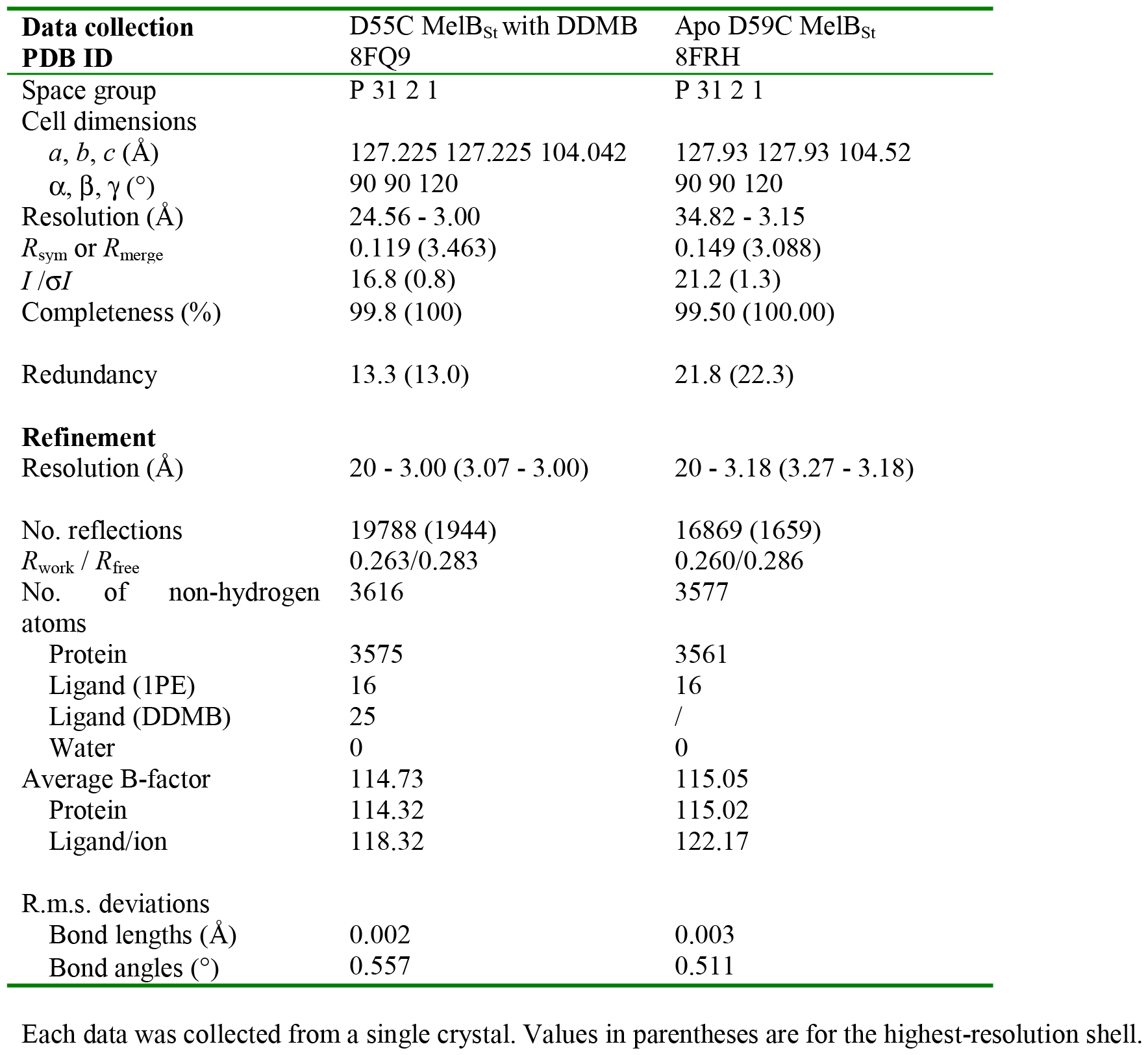
Crystallographic Data collection and refinement statistics.

The ITC measurement was also used to measure the binding of the hydrophobic ligand α-NPG to the mutant and the estimated *K*_d_ value is 15.06 ± 1.86 μM, which is significantly lower than that of WT in the absence of Na^+^ (**Table 1; Fig. 3d**). Consisting to the melibiose binding, the sugar affinity for the D55C mutant did not increase in the presence of Na^+^.

### pH effects on the H^+^-coupled melibiose transport and binding

To understand how external pH affected the H^+^-coupled symport activity, melibiose transport was carried out in 8 extracellular pH values at an interval of 0.5 units (5.5, 6.0, 6.5, 7.0, 7.5, 8.0, 8.5, or 9.0 (**Fig. 4a**). With the WT, at the acidic pH 5.5, the cells accumulated melibiose poorly, but the uptake gradually increased along with an increase in external pH values. The transport activity remained undistinguishable between external pH 7.5 – 9.0. With the D55C mutant, the external pH effect on the transport activity exhibited a pattern similar to the WT, i.e., weak activity at the acidic pH range and greater at the alkaline pH range. The steady-state accumulation level is apparently slightly higher than that in the WT. The D59C mutant showed no active transportive activity at all testing pH values, which is indistinguishable from the control cells with no MelB.

**Fig. 4.**
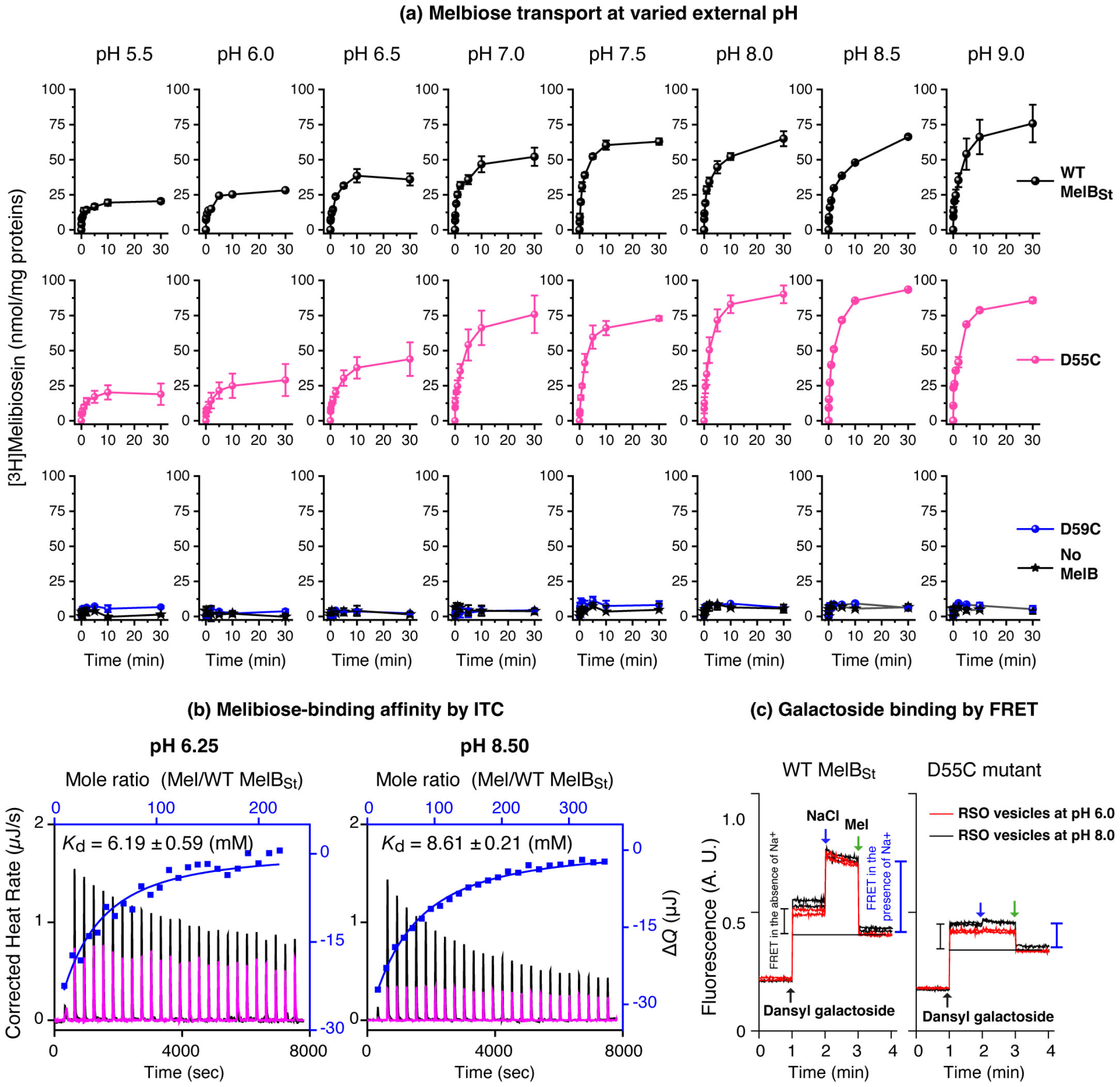
pH effects on melibiose transport and binding. **(a)** External pH effect of melibiose transport. Protein expression of the WT MelB_St_, D55C, and D59C mutants as described in **Figure 2**. After the third wash with 50-mL buffer of 100 mM KP_i_, pH 7.5, the cells were aliquoted into 8 groups and further washed with 50-mL KP_i_ buffer at pH value adjusted with from 5.5 to 9.0 at an interval of 0.5 value 0.5 and 10 mM MgSO_4_. The cell pellets collected from centrifugation were resuspended in the same buffer at a given pH value. The transport assay was conducted at 0.4 mM melibiose (specific activity, 10 mCi/mmol) in the absence of Na^+^ and Li^+^. **(b)** Melibiose binding by ITC at pH 6.25 or 8.5. The WT MelB_St_ (80 μM) in 20 mM Tris-HCl, 100 mM choline chloride, 10% glycerol, and 0.035% UDM at pH either 6.25 or 8.5 were titrated with the buffer-matched melibiose solutions at 50 mM or 80 mM, respectively. The control experiments were conducted with the same buffer solutions devoid of MelB_St_ protein. **(c)** Galactoside binding with RSO membrane vesicles. The WT or D55C MelB_St_ RSO vesicles were pre-equilibrated in 100 mM KPi buffer at pH 6.0 or 8.0 and used to measure the Trp→D^2^G FRET by sequentially adding 10 μM D^2^G, 50 mM NaCl, and >120 mM melibiose, as indicated by the arrows. The changes by the purple or blue bar indicated the FRET intensity from bound D^2^G in the absence or presence of Na^+^.

Since the transport activity was affected by the external pH value, the pH effect on sugar-binding was conducted at an acidic and an alkaline pH with two different methods. The ITC measurements of melibiose binding with the WT MelB_St_ were used to determine the binding affinity with purified protein in a Na^+^-free buffer at pH 6.25 and 8.50 as described in the method. The obtained *K*_d_ values at pH 6.25 and 8.5 are 6.19 ± 0.59 and 8.50 ± 0.21 mM, respectively. The *K*_d_ value at acidic pH was apparently slightly better at a less than a 2-fold effect, which is consistent with the sugar effect on the protonation of MelB_St_, where the absolute *K*_D(H+)_ value was from 0.56 μM to 0.26 μM in the presence of melibiose(13). This pH effect on melibiose binding result provided the missing data on the thermodynamic cycle between sugar and H^+^ as we did for Na^+^ or Li^+^ in this protein. All results support the positive cooperativity between the symport solutes.

The right-side-out membrane (RSO) vesicles expressed the WT and D55C mutants were prepared in 100 mM KPi buffer adjusted to pH 6.0 or 8.0, respectively, and the mentioned well-established Trp to dansyl galactoside FRET assay was used to detect the dansyl-galactoside binding **(Fig. 3b)**. The intensity changes in either pH or the WT and D55C mutant were similar, supporting the conclusion from the ITC measurements that the melibiose-binding affinity are less affected by the pH, which is largely different from the effect by Na^+^ or Li^+^ as reported (6, 13). Thus, the reduced transport activities observed in both WT and the D55C mutants do not stem from decreased sugar binding affinity.

### Crystal structure determination

As reported, the purified D55C and D59C MelB_St_ in detergent Detergents undecyl-β-D-maltopyranoside (UDM) solution exhibited improved thermostabilities (13). The crystal structures of D59C mutant bound with α-NPG and DDMB have been published (8). Here we report the structures of apo D59C and DDMB-bound D55C MelB_St_ mutants, which were determined by molecular replacement, modeled from positions 2 to 454 or 2 to 256 without a gap (**Fig. 5a-b**), refined to a resolution of 3.0□Å and 3.18□Å, respectively. The two structures exhibit an RMSD value of 0.263 Å. Both structures are also virtually identical to the published DDMB-bound [PDB ID 7L16] or α-NPG-bound D59C (PDB ID 7L17) structures at RMSD values of less than 0.4 Å. Notable, both mutants do not bind Na^+^ or Li^+^ nor catalyze Na^+^- or Li^+^-coupled sugar transport (**Figs. 2-4**). In both structures, a PEG molecule was modeled between the helices IX and XII at the cytoplasmic side, which could imply a potential lipid-binding site.

**Fig. 5.**
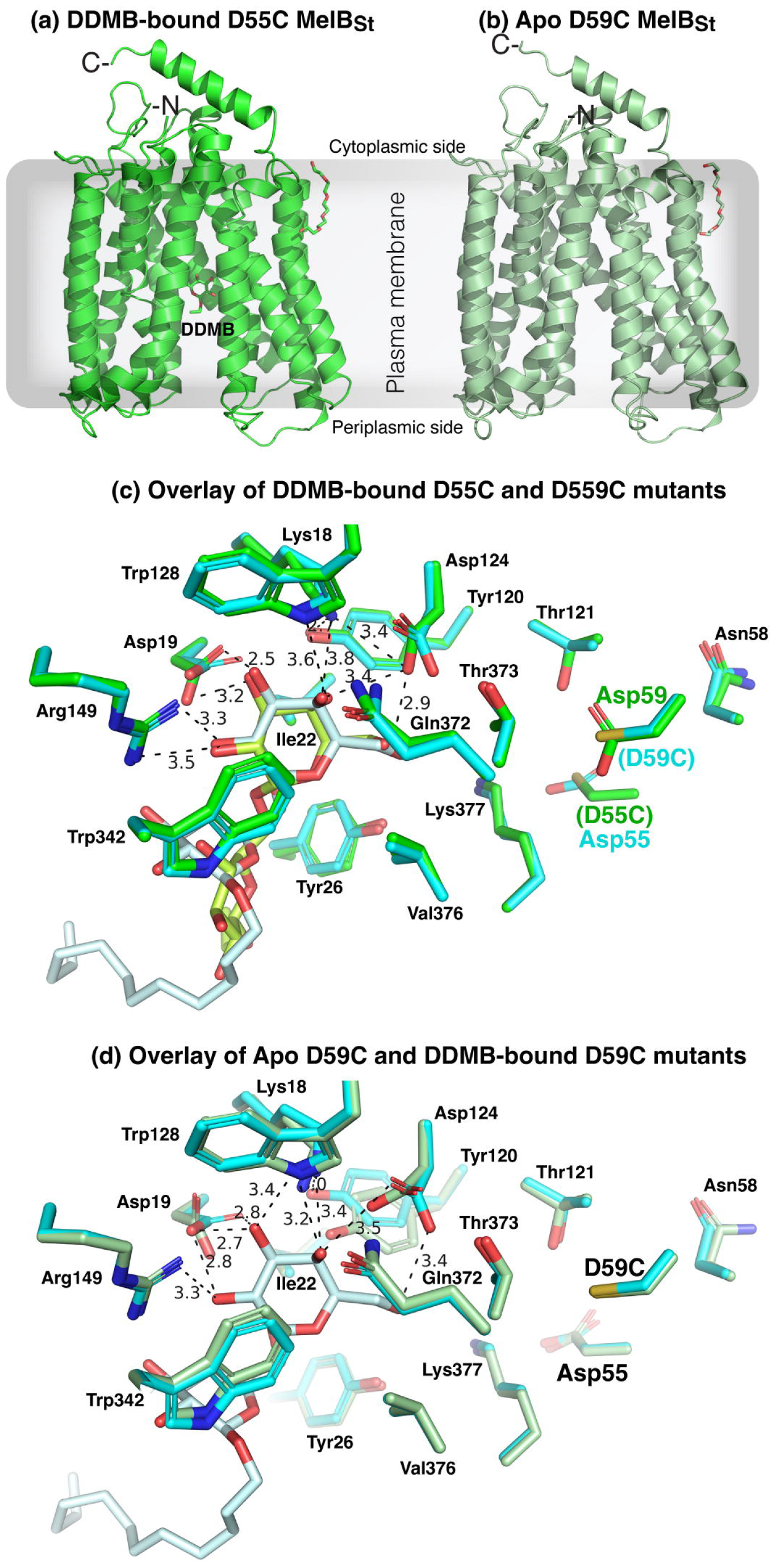
Crystal structures of the D55C and D59C MelB_St_ mutants. **(a)** D55C mutant bound with DDMB refined to a resolution of 3.0 Å. The structure was shown in cartoon representation in green and DDMB with 2-carbon tail is shown in the stick and labeled. **(b)** D59C mutant at apo state refined to a resolution of 3.18 Å. The structure was shown in cartoon representation in light green. For both structures, the helices V and VIII were placed at the middle and N-terminus was placed at the left side. One bound PEG molecule interacted with the cytoplasmic side of helix IX in both structures. The sidedness of the membrane is indicated. **(c)** The specificity determinant pockets for the sugar substrates and the coupling cations. The DDMB-bound D55C mutant was overlaid on the DDMB-bound D59C mutant [PDB 7L16]. The DDMB-binding residues are shown in the sticks, and the DDMB molecules were colored green (D55C mutant) and pale cyan (D59C mutant), respectively. The cation-binding residues (positions 55, 58, 59, and 121) were shown in the stick. **(d)** Overlay of the apo D59C and DDM-bound D59C mutants. The DDMB-binding residues are shown in the sticks, and the DDMB molecules were colored green (D55C mutant) and pale cyan (D59C mutant), respectively. The cation-binding residues (Asp55, Ans58, D59C, and Thr121) were shown in the stick.

Apo D59C and D55C structures showed virtually identical galactoside-binding pockets (**Fig. 5c**). In the D55C structure, the galactosyl moiety of bound DDMB exhibits a similar pose to that in the D59C bound with either DDMB or α-NPG. The densities of the 11-carbon tail were disordered and only modeled 2 carbons. The glucosyl moiety poses different from that in the D59C structure. The Na^+^-binding pocket in both crystal structures and the previously published D59C structures are nearly identical with a Cys side chain present at positions either Asp55 or Asp59, respectively (**Fig. 5c-d**).

In comparison with the Na^+^-bound WT MelB_St_ cryoEM structure (**Fig. 6**), both the DDMB-bound D55C and apo D59C MelB_St_ mutants exhibited loosely packed side chains of this binding pocket due to the absence of Na^+^ and the D55C structure was removed from clarity.

**Fig. 6.**
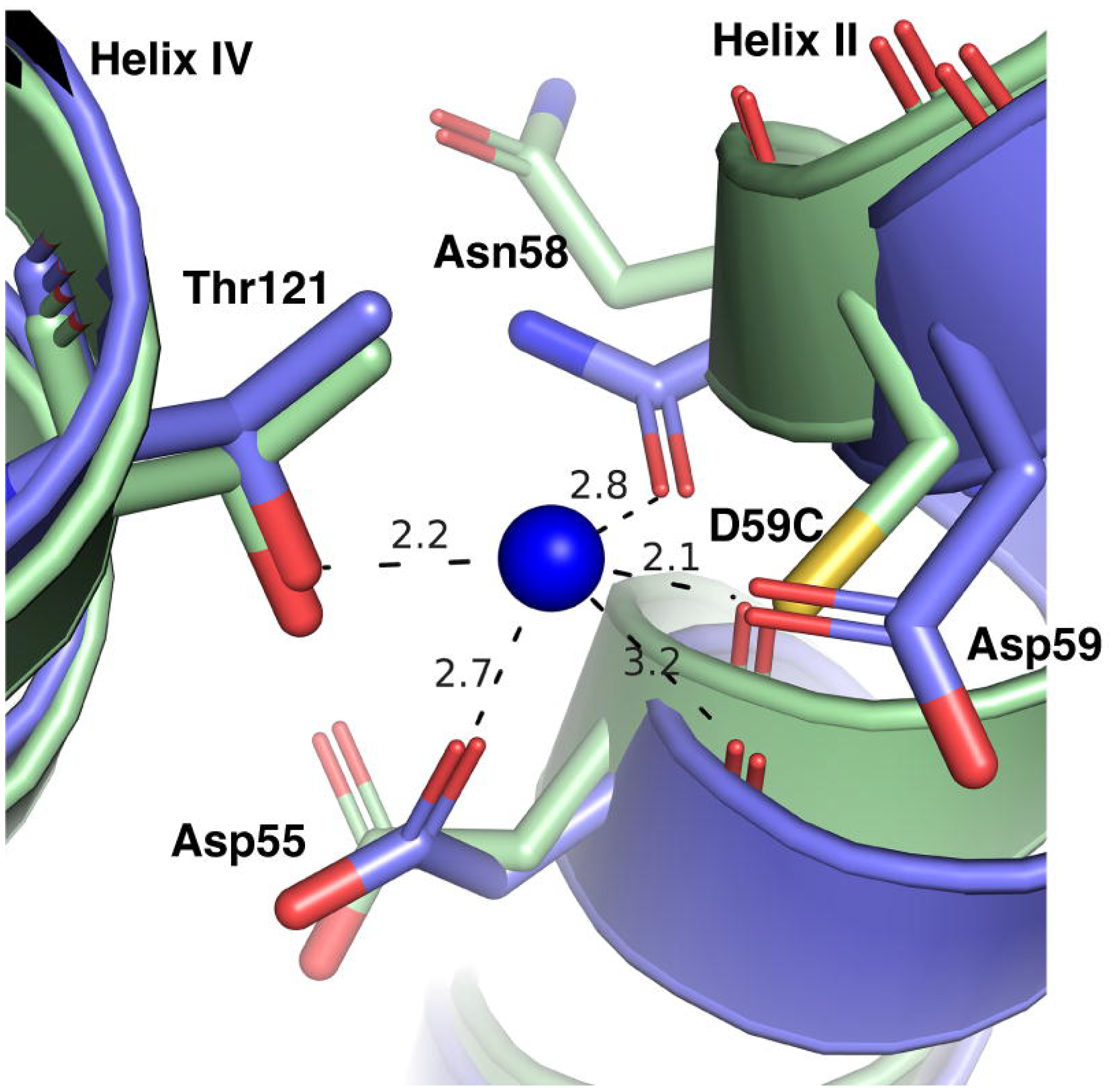
Superimposed Na^+^-binding pocket of the WT and mutants. The N-terminal helices of the Na^+^-bound inward-facing cryoEM structure of the WT MelB_St_ [PDB ID 8T60] were aligned on the Apo D59C mutant. The Na^+^-binding residues were shown in the stick, and the distance of Na^+^ coordinates was indicated with dashed lines (Å). Helices II and IV were labeled.

### Effect of single-site mutations on Na^+^-binding affinity

The three well-characterized single-site mutants D55C, D59C, or T121A were subjected to free energy calculations to quantitatively analyze the effects of mutation on the Na^+^ binding affinity. The change in the binding affinity upon mutation was evaluated by the free-energy perturbation (FEP) simulations (**Fig. 7a; Table 3, supplementary information Table S1**). The thermodynamics cycle for Na^+^ binding was constructed. The Δ*G*_(Na+-unbound)_ reflects the free energy change upon mutating the WT to mutant in the Na^+^-unbound state of MelB_St_. The Δ*G*_(Na+-bound)_ reflects the free energy change upon mutating the WT to the mutant at the Na^+^-bound state. Based on the thermodynamic cycle, the difference between these two free energy changes (ΔΔ*G*_Na+ binding =_ Δ*G*_mutation, Na+-bound state_ - Δ*G*_mutation, Na+-unbound state_) equals the change in the Na^+^-binding free energy induced by the mutation. In other words, ΔΔ*G*_Na+ binding_ reflects how much the mutation perturbs the binding affinity of Na^+^. A more positive ΔΔ*G* value indicates a greater reduction in cation-binding affinity due to the mutation. The D59C mutation yielded the greatest destabilization in Na^+^ binding with a ΔΔ*G* value of 10.5 ± 0.3 kcal/mol, followed by the D55C mutation with a ΔΔ*G* value of 8.6 ± 0.3 kcal/mol, and the T121A mutation with a least ΔΔ*G* value of 6.7 ± 0.1 kcal/mol. The data are consistent with the notion that all side chains are important but the two carboxyl groups on Asp55 and Asp59 play major roles in the Na^+^ binding in MelB_St_.

**Table 3.** Mutation-induced free energy changes of Na^+^ or H^+^ binding (ΔΔ*G*_cation binding_)

| Quantity | WT → D59C<br>(kcal/mol) | WT → D55C<br>(kcal/mol) | WT → T121A<br>(kcal/mol) |
| --- | --- | --- | --- |
| $\Delta G_{\text{mutation, Na}^+-\text{unbound}}^*$ | 117.2 ± 0.2 | 124.3 ± 0.2 | 15.89 ± 0.03 |
| $\Delta G_{\text{mutation, Na}^+-\text{bound}}^*$ | 127.7 ± 0.1 | 132.9 ± 0.2 | 22.5 ± 0.1 |
| $\Delta\Delta G_{\text{Na}^+ \text{ binding}}^{***}$ | **10.5 ± 0.3** | **8.6 ± 0.4** | **6.7 ± 0.1** |
| $\Delta G_{\text{mutation, Asp59}}^{-*}$ | | 124.2 ± 0.1 | |
| $\Delta G_{\text{mutation, Asp59}}^{-H**}$ | | 126.2 ± 0.1 | |
| $\Delta\Delta G_{\text{proton binding at Asp59}}$ | | **2.0 ± 0.2** | |
| $\Delta G_{\text{mutation, Asp55}}^{-*}$ | 122.2 ± 0.3 | | |
| $\Delta G_{\text{mutation, Asp55}}^{-H**}$ | 117.2 ± 0.2 | | |
| $\Delta\Delta G_{\text{proton binding at Asp55}}^{***}$ | **5.1 ± 0.4** | | |

<sup>\*</sup>, $\Delta G_{\text{mutation, cation-unbound state}}$ reflects the free energy change induced by a mutation when MelB<sub>St</sub> is at a cation-free form or unbound with the test cation (Na<sup>+</sup> or H<sup>+</sup>).
<sup>\*\*</sup>, $\Delta G_{\text{mutation, cation-bound state}}$ reflects the free energy change induced by mutation when MelB is bound with the cation.
<sup>\*\*\*</sup>, $\Delta\Delta G_{\text{cation-binding}} = \Delta G_{\text{mutation, cation-bound state}} - \Delta G_{\text{mutation, cation-unbound state}}$ . $\Delta\Delta G_{\text{cation-binding}}$ . It reflects the change in the cation-binding free energy induced by a mutation. A more positive $\Delta\Delta G$ indicates a larger reduction in cation-binding affinity after the mutation.

**Fig. 7.**
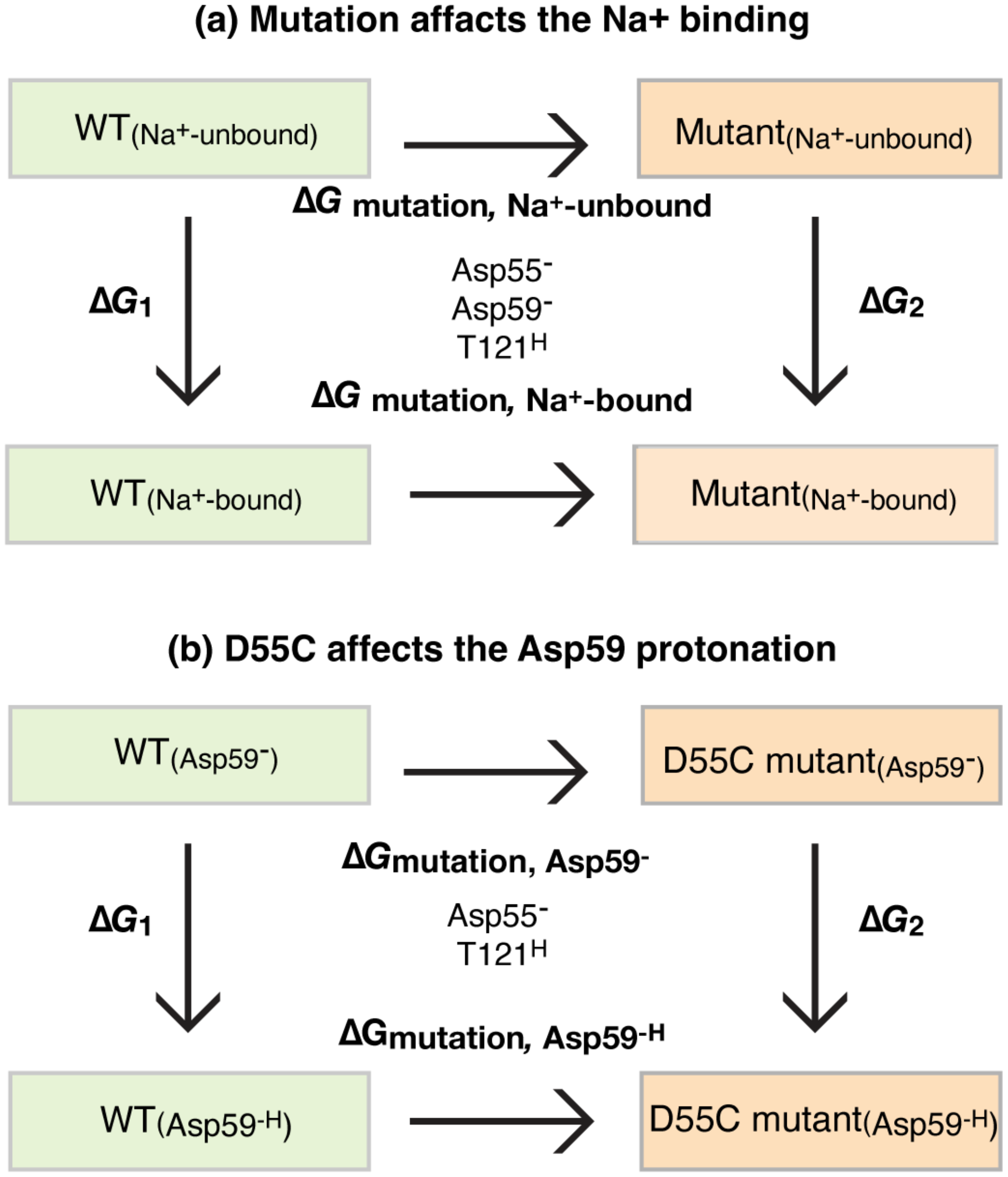
Thermodynamic cycle. The simulations were carried out as described in the Methods and data were presented in **Table 3**.

### The effect of mutations on the pronation of Asp55 or Asp59

Among the two titratable residues, a largely elevated p*K*a value of 8.75 for Asp59 was estimated by PROPKA (14). PROPKA estimation on Asp55 obtained from a consistent simulation protocol reveals a value of 4.0, in the expected range for an Asp side chain in bulk solution, which further solidifies that Asp59 is the sole protonation site in MelB. To determine how the Cys mutation at Asp55 affects the protonation of Asp59 and vice versa, free-energy perturbation simulations were applied to calculate the ΔΔ*G*_proton binding at Asp59_ (**Table 3**). Based on the thermodynamic cycle, this ΔΔ*G* equals the difference between the mutation-introduced free energy change in the H^+^-unbound and H^+^-bound states of MelB_St_, with the protein binding site being either Asp59 or Asp55 (**Fig. 7b; Table 3**). Thus, this ΔΔ*G* value reflects the change of the proton binding affinity of either residue upon the mutation of the other. A more positive value indicates a weaker proton binding affinity or larger inhibition of the proton binding introduced by mutation. The FEP simulation results showed that the H^+^-binding affinity of Asp59 in the D55C mutant is only 2.0 ± 0.2 kcal/mol lower than the WT, i.e., the p*K*a value is estimated to decrease by 1.5 units. Compared to the effect on the Na^+^ binding, the effect of the D55C mutation on the proton binding of Asp59 is much smaller (**Table 3**). In contrast, the D59C mutation leads to a decrease of the H^+^-binding affinity of Asp55 by 5.1 ± 0.4 kcal/mol, about a 3.7-unit decrease in the p*K*a value. The data support that the D55C mutant remains the H^+^-coupled symport activity mediated by the protonation and deprotonation of the Asp59 residue (**Fig. 6)**. In contrast, the D59C mutation eliminates this activity due to the removal of the sole proton binding site (Asp59), and the remaining Asp55 is incapable of serving as a protonation site because its proton-binding affinity is further decreased by the Cys mutation at Asp59.

## Discussion

There are two well-conserved acidic residues in the cation-binding pocket of MelB, and those two positions have also been well-studied in the past by several laboratories using varied methods (5, 7, 8, 17, 19, 20). Cys or Ala mutants on the strictly conserved Asp59 lose the binding of Na^+^ and Li^+^, as well as the melibiose active transport coupled to the Na^+^, Li^+^, or H^+^. However, these mutants still mediate melibiose translocation driven by melibiose concentration gradient (**Fig. 2a, inset**), converting the WT MelB_St_ symporter to a uniporter (8).

The D55C mutant also loses the Na^+^ and Li^+^ binding and the melibiose active transport coupled to the Na^+^ or Li^+^, but maintained the melibiose concentration gradient-drive melibiose transport as shown by the melibiose fermentation assay (**Fig. 2a inset**), similar to the D59C mutant. Interestingly, the single-Cys D55C mutant at a Cys-less background with endogenous 8 Cys residues replaced with Ala, showed poor melibiose fermentation activity (19). The protein stability of the Cys-less MelB_St_ is not as good as the WT, which might result in poor melibiose transport and fermentation. The D55C MelB_St_ retains the H^+^-coupled symport activity, and the uncoupler CCCP abolished both activities (**Fig. 2b**). The crystal structures of both D55C and D59C mutants in the absence or presence of a ligand reveal a virtually identical cation-binding pocket except for the mutations and sidechain poses of major Na^+^-binding players (**Figs. 5-6**).

To further characterize the individual role of the three well-studied side chains (Asp 55, Asp59, and Thr121) in this shared cation-binding pocket, the FEP simulations were utilized to estimate the mutational effects of these residues on the cation-binding affinities for Na^+^ and H^+^ in MelB_St_. The data showed that all three side chains are important for the Na^+^ binding and the single-site mutations at the two negatively charged Asp59 and Asp55 residues make greater destabilization for the Na^+^ binding, which supports the experimentally established conclusion that the Na^+^ binding is primarily stabilized by the carboxyl groups in the side chains of Asp55 and Asp59, especially Asp59 (7, 11, 13).

PROPKA calculations suggest that the estimated p*K*a values of Asp59 and Asp55 are 8.7(14) and 4.0, respectively. The highly elevated p*K*a value of Asp59 is consistent with its major role as the proton-binding site(8), and the normal range of p*K*a value obtained at the position Asp55 also supported that Asp55 is not involved in the protonation of the WT MelB_St_. When Thr121 is mutated by Ala, proton-binding affinity at Asp59 is unchanged (14). The free energy calculation (**Fig. 7b**) showed that the proton-binding affinity of Asp59 is only slightly reduced by 2.0 ± 0.2 kcal/mol by D55C mutation, largely retaining Asp59’s capability of binding protons. In contrast, Asp55’s binding affinity is reduced by 5.0 ± 0.4 kcal/mol by D59C mutation, leaving no proton-binding site in the D59C mutant. This computational result is consistent with the experimental observation that the D59C mutation effectively eliminates proton-coupled transport activity while the D55C mutation selectively eliminates the Na^+^ and Li^+^ binding and converts MelB_St_ to a solely H^+^-coupled symporter.

We have reported that the sugar-binding affinity is conformation-sensitive, but both the inward- and outward-facing conformations of MelB_St_ exhibited similar cation-binding pockets(8, 11, 14). This has raised an important question about the cation-releasing mechanisms during sugar/cation symport cycling. It is interesting that Na^+^ binding in MelB can be selectively eliminated readily by a single-site mutation on either Asp55 or Thr121 position without significant effect on H^+^ in both positions or on Li^+^ binding in the Thr121 position, as well as coupled transport (8, 14). It is likely that the Na^+^ coordinates required stringent positioning of each sidechain. We are attempting to explain that the Asp55 side chain rotamer movement after releasing the sugar could release the Na^+^ to the cavity. Further, the Na^+^-binding affinity is greater than the intracellular Na^+^ concentrations, implying that the membrane potential (ΔΨ, inside negative) is needed for releasing the Na^+^ to the cytoplasm. This is consistent with the previous study showing that ΔΨ is required for Na^+^-coupled melibiose uptake(4).

The D55C MelB_St_ with only one H^+^-coupling mode is a useful tool to study the H^+^-coupled transport without interference by potentially contaminated Na^+^. When conducting the H^+^-coupled transport at external acidic pH values (pH< 7.0), it is surprising that both WT and the D55C mutant MelB_St_ exhibited largely reduced transport activities. The binding assays using two methods with purified proteins and RSO membrane vesicles showed that the sugar-binding affinity at both acidic and alkaline pH is reasonably well-preserved, which excluded the possibility of poor sugar-binding affinity at acidic pH values. Normally, *E. coli* cells maintain a relatively constant intracellular pH value of 7.6(21, 22). Even at the extracellular pH of 5.5 and 9.0, the intracellular pH was determined as 7.4 and 7.8, respectively (21). Thus, the decrease in the external pH should increase the pH gradient between external and internal (ΔpH, inside alkaline). If the transport rate were coupled to bulk-liquid ΔpH (inside alkaline), greater transport activities should have been expected at lower external pH values. In contrast, the data revealed that the greater the ΔpH value (such as external liquid pH 5.5), the poorer the transport activity, which does not support the ΔpH as a major driving force for the H^+^-coupled melibiose uptake mediated by MelB_St_. On the other hand, greater transport activities were observed at reversal ΔpH conditions (external liquid pH>=8.0). All the data support that the H^+^/melibiose symport is mainly coupled to ΔΨ.

It is well known that an H^+^-coupled secondary active transport is coupled to the total free energy as a form of the electrochemical H^+^ gradient 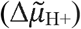, also named proton motive force (23, 24). The 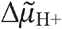 at least consists of two compensating terms: ΔΨ and ΔpH. It is also known that the increase in ΔpH resulted in a decrease in the ΔΨ value, and vice versa (25). The poor transport activities obtained at external acidic pH values or at greater ΔpH (internal alkaline) could stem from the reduced ΔΨ values.

Another unexpected result is the CCCP-sensitive strong transport activity in the alkaline pH range (pH>8.0) with both WT and the D55C mutant. The experimentally determined p*K*a value of the cation pocket, i.e., the Asp59, is approximately 6.5 with purified MelB_St_(13). The previous buffer protonation study by measuring enthalpic change at different buffer systems reveals that at pH 8.2, nearly all MelB_St_ is deprotonated(13). The possibility that this phenomenon arises from Na^+^ contaminations can be undoubtedly excluded since this D55C mutant does not bind Na^+^. Thus, the observed H^+^-coupled active melibiose transport at low external H^+^ concentration or at revered pH gradients (inside, relatively acidic) indicated that the WT and D55C mutant MelB_St_ carried by the live-intact cells can be protonated during the transport cycle even at an external pH 9.0 value. The data support a recent novel theory over Peter Mitchell’s chemiosmosis theory on the electrochemical H^+^ gradient 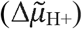, where a third term named transmembrane-electrically localized protons (TELP) was described (26, 27), in addition to ΔΨ and ΔpH. Accordingly, the transmembrane-electrically localized protons density is likely several magnitudes higher than the bulk solution H^+^ concentration because the plasma membrane acts as a TELP-associated capacitor, where the intracellular negative charges (OH^−^) transmembrane-electrostatically attract trap H^+^ onto the periplasmic surface of the plasma membrane. This revision of the chemiosmosis theory can also explain the transport bioenergetics of the alkalophilic bacteria (27).

Together with the previous studies, the structural and functional analysis, as well as the quantification of the energetic contributions of the individual residues in the MelB cation-binding pocket, clearly demonstrate that Asp55, Asp59, and Thr121 are critical for Na^+^ binding, Asp55 and Asp59 are critical for Li^+^ binding, and Asp59 is the sole H^+^-binding site. Furthermore, our study also determined the membrane potential as the primary driving force for the H^+^-coupled active melibiose transport mediated by MelB.

## Experimental Procedures

### Reagents

[1-^3^H]Melibiose (5.32 Ci/mmol) was custom synthesized by *PerkinElmer*, and unlabeled melibiose was purchased from *Acros Organics* (*Fisher Scientific*). 2′-(*N*-dansyl)aminoalkyl-1-thio-β-D-galactopyranoside (D^2^G) was gifted from Dr. Gerard Leblanc. MacConkey agar media (lactose-free) was purchased from *Difco*. Detergents undecyl-β-D-maltopyranoside (UDM), and octyl-β-D-glucoside (OG) were purchased from *Anatrace*. Detergent dodecyl-β-D-melibioside (DDMB) and *E. coli* lipids (Extract Polar, 100600) were purchased from *Avanti Polar Lipids, Inc*. All other materials were reagent grade and obtained from commercial sources.

### Strains and plasmids construction

*E. coli* DW2 cells (*mel*A^+^, *melB*^−^, *lacZ*^−^*Y*^−^) (28) were used for protein expression and functional studies. The expression plasmid pK95 ΔAH/MelB_St_/CHis_10_ was used for constitutively expressing the WT MelB_St_ and MelB_St_ mutant D55C and D59C.

### [1-^3^H]Melibiose transport assay

*E. coli* DW2 cells transformed with a given plasmid were grown in Luria-Bertani (LB) broth with 100 mg/L ampicillin in a 37 °C shaker. The overnight cultures were diluted by 5% to fresh LB broth with 0.5% glycerol and 100 mg/L ampicillin and shaken at 30 °C for 5 h. The cells were washed with 50-ml 100 mM KP_i_, pH 7.5 three times, followed by washing with the assay buffer (100 mM KP_i_, pH 7.5, 10 mM MgSO_4_). The cell pellets were resuspended and adjusted to A_420_ = 10 (∼0.7 mg proteins/mL) with the assay buffer. The transport assay was performed at 0.4 mM [^3^H]melibiose (specific activity of 10 mCi/mmol) in the absence of Na^+^ and Li^+^ or the presence of Na^+^ or Li^+^ as described (4, 29) to test the three transport model coupling to H^+^, Na^+^, or Li^+^, respectively. The transport time courses were carried out by quenching the cells at zero, 5 s, 10 s, 30 s, 1 m, 2 m, 5 m, 10 m, and 30 m by dilution and fast filtration. The filters were subjected to radioactivity measurements using a liquid scintillation counter.

The CCCP effect on the transport activity carried out in the absence of Na^+^ and Li^+^ was conducted by incubating the extensively washed Na^+^-free cells with 10 μM CCCP for 10 min prior to the transport assay.

### pH effect on transport activity

*E. coli* DW2 cells expressing the WT MelB_St_, D55C, or D59C mutants were prepared as described above. After three washing using the Na^+^-free buffer, the last washing and resuspending were performed using a specific pH-adjusted buffer. By altering the ratio of KH_2_PO_4_ and K_2_HPO_4_, the buffers were adjusted to 5.5 – 9.0 with an interval of 0.5 value. A small amount of Tris base was added to prepare the pH 9.0 solution.

### Melibiose fermentation on MacConkey agar plates

*E. coli* DW2 cells were transformed with a plasmid carrying the WT or mutants and plated on MacConkey agar plates containing 30 mM melibiose, 100 mg/L of ampicillin, and incubated at 37 °C. After 18 h, the plates were viewed and photographed immediately. Magenta color colonies, a normal melibiose fermentation; yellow colonies, no melibiose fermentation due to poor transport.

### MelB_St_ protein expression and purification

Cell growth for the large-scale production of WT MelB_St_ was carried out using the same expression vector and cell strain as used in the functional analyses (7, 30). Briefly, MelB_St_ purification from membranes by cobalt-affinity chromatography (Talon Superflow Metal Affinity Resin, Takara) after extraction by 1.5% UDM. MelB_St_ protein was eluted with 250 mM imidazole in a buffer containing 50 mM NaPi, pH 7.5, 200 mM NaCl, 0.035% UDM, and 10% glycerol, and further dialyzed to change the buffer conditions accordingly.

### Protein concentration assay

The Micro BCA Protein Assay (Pierce Biotechnology, Inc.) was used to determine the protein concentration.

### Isothermal titration calorimetry

All ITC ligand-binding assays were performed with the TA Instruments (Nano-ITC device) as described (13), which yields the exothermic binding as a positive peak. In a typical experiment, the titrand (MelB_St_) placed in the ITC Sample Cell was titrated with the specified titrant Na^+^, Li^+^, melibiose, or α-NPG (placed in the Syringe) in the assay buffer by an incremental injection of 2-μL aliquots at an interval of 300 sec at a constant stirring rate of 250 rpm (nano-ITC). MelB_St_ protein samples were buffer-matched to the assay buffer by dialysis. For measuring Na^+^ or Li^+^ binding, 50 mM choline chloride was supplemented into the Na^+^-free buffer.

All samples were degassed using a TA Instruments Degassing Station (model 6326) for 15 min. Heat changes were collected at 25 °C, and data processing was performed with the NanoAnalyze (version 3.7.5 software) provided with the instrument. The normalized heat changes were subtracted from the heat of dilution elicited by the last few injections, where no further binding occurred, and the corrected heat changes were plotted against the mole ratio of titrant versus titrand. The values for the binding association constant (*K*_a_) were obtained by fitting the data using the one-site independent-binding model included in the NanoAnalyze software (version 3.7.5). The dissociation constant (*K*_d_) = 1/*K*_a_.

### Preparation of RSO vesicles

Right-Side-Out (RSO) membrane vesicles were prepared from *E. coli* DW2 cells overexpressing the WT or D55C MelB_St_ by osmotic lysis (4, 31, 32). The concentrated RSO membrane vesicles at protein concentrations of 20 mg/ml in a 100 mM KP_i_ buffer (pH 7.5) were diluted to a protein concentration of 1 mg/ml using 100 mM KPi at pH 6.0 or pH 8.0, respectively. The sample were incubated at 23 °C for 30 min prior to binding assay.

### Galactoside-binding assay

Duplicate measurements of Trp→Dansyl galactoside (D^2^G) FRET experiments were conducted using an Amico-Bowman Series 2 (AB2) Spectrofluorometer (4). An aliquot of 200 μL RSO vesicles at pH 6.0 or 8.0 was used to monitor the fluorescent changes. Trp residues were excited at 290 nm and emission of D^2^G was recorded at 490 nm. On the time trace, the sequential additions of 10 μM D^2^G, 50 mM NaCl, and >120 mM melibiose were carried out at 1-, 2- and 3-min time points, respectively.

### Crystallization, native diffraction data collection, and processing

The purified D55C or D59C MelB_St_ mutant in solutions were dialyzed overnight against the sugar-free dialysis buffer (consisting of 20 mM Tris-HCl, pH 7.5, 100 mM NaCl, 0.035% UDM, and 10% glycerol), concentrated with Vivaspin column at 50 kDa cutoff to approximately 30-50 mg/ml, and subjected to ultracentrifugation at 384,492□*g* for 45□min at 4□°C (Beckman Coulter Optima MAX, TLA-100 rotor), stored at −80 °C after flash-frozen with liquid nitrogen for crystallization trials. A phospholipid stock solution at a concentration of approximate 20 mM were prepared by dissolving the *E. coli* Extract Polar (Avanti, 100600) with a dialysis buffer containing 0.01% DDM instead of 0.035% UDM.

Crystallization trials were carried out by the hanging-drop vapor-diffusion method at 23□°C by mixing 2-μL pre-treated protein samples with 2-μl reservoir. For the crystallization of the apo D59C MelB_St_ mutant, the protein sample was diluted to a final concentration of 10 mg/ml with the same sugar-free dialysis buffer, supplemented with phospholipids at a concentration of approximately 3.6□mM from the 20-mM stock, and incubated for 15 min prior to the crystallization trials. The apo D59C mutant crystals appeared in 3-4 days against a reservoir consisting of 100 mM Tris-HCl, pH 8.5, 100 mM NaCl_2_, 50 mM CaCl_2_, and 32% PEG 400, and frozen with liquid nitrogen in 2 weeks, and tested for X-ray diffraction at the Lawrence Berkeley National Laboratory ALS BL 8.2.2 via remote data collection method.

For the crystallization of the DDMB-bound D55C MelB_St_ mutant, the protein solution was diluted with the dialysis buffer to a final concentration of 10 mg/ml with the supplement of phospholipids at a concentration of approximate 3.6□mM from the 20-mM stock and 0.015% DDMB (1 x CMC) and 10% PEG 3350 and incubated for 15 min prior to the preparation of the crystallization drops. The crystals were grown using a reservoir consisting of MES, pH 6.5, 100 mM NaCl_2_, 50 mM CaCl_2_, 100 mM and 32% PEG 400. Crystals appeared in 3-4 days, were frozen with liquid nitrogen in 3 weeks, and tested for X-ray diffraction at ALS BL 5.0.1 via remote data collection method. The complete diffraction datasets for the apo D59C and DDMB-bound D55C structures were collected at 100□K from a single cryo-cooled crystal at a wavelength of 1.0 Å on ALS BL 8.2.2 with a ADSC Quantum 315r detector or 0.97741□Å on ALS BL 5.0.1 with a Dectris Pilatus 2M detector, respectively. ALS auto-processing XDS or DIALS programs output files were further reduction by AIMLESS in the ccp4i2 program for the structure solution (33). The statistics in data collection are described in **Table 3**.

### Structure determination

The structure determination for the apo D59C and DDMB-bound D55C MelB_St_ mutants was performed by molecular replacement method using the α-NPG-bound D59C MelB_St_ mutant structure [PDB ID 7L17] as the search template, followed by rounds of manual building and refinement to resolutions of 3.18 Å [PDB ID 8FRH] or 3.0 Å [PDB ID 8QF9], respectively, using Phenix(34). The 3-D structure of DDMB (code, LMO) from the DDMB-bound D59C mutant [PDB ID 7L16] was used to fit and 10 carbon atoms were removed due to disorder. The apo D59C and DDMB-bound D55C structures were modeled from positions 2 to 454 or 2 to 456, respectively, without a gap. In both density maps, there was a strong positive density with a sausage shape aligning with the helix IX. A PEG molecule (ligand ID 1PE) was used to model for both structures. All structures are virtually identical.

There is an unknown denticity blob in the sugar-binding pocket of the apo D59C structure, which is too small to fit even by a monosaccharide. It might be fit by a glycerol molecule. The statistics of refinement for the final models are summarized in **Table 3**. For apo and DDMB-bound structures: Ramachandran favored of 95.57% and 94.04%, Ramachandran outliers of 0% and 0.44%, clash scores of 1.38 and 2.18, and overall scores of 1.18 and 1.55, respectively, as judged by MolProbity in Phenix. Pymol (2.5.2) was used to generate all graphs(35).

### MD simulations

The initial configuration for our simulations was derived from the crystal structure of the D55C mutant of MelB_St_. For WT system setup, the Asp55 was reverted to a deprotonated state. For the D55C mutant setup, the protein structure was used directly. For the D59C mutant setup, the Asp59 was reverted to a deprotonated state, and the Asp55 was mutated to Cys. For the T121A mutant setup, the Asp59 was deprotonated, and the Thr121 was mutated to Ala. For all of the systems, the Asp59 and Asp55 residues were deprotonated whenever applicable, while other ionizable residues maintained their standard protonation states (deprotonated Asp, Glu, His, and protonated Thr, Ser, Cys, Lys, Arg, Tyr). In the initial setups of all three titratable systems, a melibiose molecule was positioned within the sugar-binding site based on the crystal structure of α-NPG, and the Na^+^ ion was positioned in the crystal structure. The protein was integrated into a lipid bilayer composed of a 7:2 ratio of POPE to POPG, mirroring the *E. coli* membrane composition, with a total of 288 lipid molecules. Water molecules enveloped both sides of the lipid bilayer, and a solution with an approximate concentration of 0.15 M was created by introducing NaCl ions among the water molecules. The simulation system encompassed approximately 121,000 atoms, and the periodic boundary condition was set at approximately 100 Å × 100 Å × 123 Å. The CharmmGUI web interface(36) was utilized to construct the system.

The system was equilibrated through the following procedure. Initially, the systems were relaxed through geometry optimization, with harmonic restraints (500 kJ/mol/Å^2^ force constant) applied to the heavy atoms of the protein and lipids. Subsequently, a 250 ps simulation at a temperature of 303.15 K was conducted in the constant NVT ensemble. The temperature of the system was maintained using a Langevin thermostat with a friction coefficient of 1 ps□^1^. The restraints were then systematically diminished to zero during 2 ns of dynamics in the constant NPT ensemble at 303.15 K and 1 atm pressure. The pressure of the system was maintained using the Langevin piston Nose-Hoover method (37, 38) piston pressure A 200 ns production trajectory was generated in the constant NPT ensemble, maintaining the same temperature and pressure., and a 2 fs timestep was employed to propagate the trajectory, with constraints on bond lengths involving hydrogen atoms. The CHARMM36 force field(39-42)was employed for the protein, lipids, and melibiose, and the TIP3P model for water molecules(43). Electrostatic interactions were computed using the particle mesh Ewald method (44) with a 12 Å cutoff for van der Waals interactions. All MD simulations were performed using the NAMD software package(45).

### FEP simulations

To assess the alteration in binding free energy resulting from mutations, we conducted free-energy perturbation (FEP) simulations for each mutant paired with the WT, namely WT⟷D59C WT⟷D55C, and WT⟷T121A. In each FEP simulation, the side chain of the mutated residue was systematically transitioned from its original state to the new one by gradually modifying the system’s Hamiltonian using a λ parameter ranging from 0 to 1. Essentially, the λ parameter serves to linearly interpolate between the Hamiltonians of the WT system and the mutant. Equation 1 describes the Hamiltonian of the system undergoing mutation.

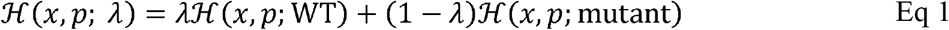

As λ progresses from 0 to 1, the system’s Hamiltonian undergoes a gradual transition from the description of one variant of MelB_St_ (e.g., WT) to another (e.g., mutant). From each FEP simulation, we extracted the free energy change (Δ*G*) for the entire system resulting from the mutation, including the cation-unbound and cation-bound states of MelB_St_, yielding Δ*G*_mutation, cation-unbound state_ and Δ*G*_mutation, cation-bound state_, respectively.

Subsequently, we construct a thermodynamic cycle (Fig. 7), allowing us to calculate the difference in the cation binding free energies (Na^+^ or proton) between the mutant and WT (Eq 1):

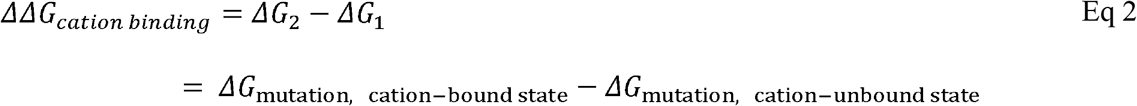

A positive ΔΔ*G* indicates a decrease in binding affinity upon mutation. In the context of proton binding affinity for a specific residue, the terms “cation-bound” and “cation-unbound” can be alternatively replaced with “protonated” and “deprotonated” states of the residue, respectively.

To reduce the statistical uncertainty in the calculated ΔΔ*G*, for every mutation pair in each condition (bound or unbound), we conducted FEP simulations in both forward and backward directions, i.e., WT → mutant and mutant → WT. This approach enables the application of the simple overlap sampling (SOS) technique for the estimation of Δ*G*_mutation, cation-bound state_ or Δ*G*_mutation, cation-unbound state_ along with their error bars. The free energy change within each window of the FEP was calculated using Eq 3. The simulations were carried out under isothermal and isobaric conditions (constant pressure and temperature, i.e., NPT ensemble).

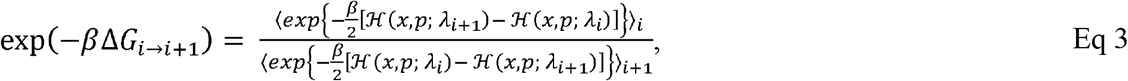

where Δ*G*_*i*…*i*+1_ *represents the Gibbs free energy change when the system transitions from the state represented with λ*= *λ*_*i*_ *to λ*= *λ*_*i*+1_, ℋ(*x, p*; *λ*_*i*+1_) - ℋ(*x, p*; *λ*_*i*_) denotes the energy difference between the two Hamiltonians with *λ* = *λ*_*i*_ and *λ* = *λ*_*i*+1_, at coordinates (x) and momenta (p) for all atoms in the system, and ⟨… ⟩ in denotes ensemble averages. In the forward run, the coordinates (*x*) and momenta (p) sampled under ℋ(*x, p; λ*_*i*_) are used to calculate ℋ(*x, p*; *λ*_*i*+1_) - ℋ(*x, p*; *λ*_*i*_) at each snapshot. In the backward run, the *x*’s and *p*’s sampled under ℋ(*x, p*; *λ*_*i*+1_) are used to calculate ℋ(*x, p*; *λ*_*i*_) - ℋ(*x, p*; *λ*_*i+1*_) at each selected snapshot. The parameter β is defined as *1/k*_*B*_*T*, where *k*_*B*_ and T denote Boltzmann’s constant and absolute temperature, respectively.

In each FEP run for the mutation process, we simulated 20 windows with λ values gradually transitioning from 0 to 1 for the forward run or 1 to 0 for the backward run, using a 0.05 interval. Within each window, we conducted 5 ns sampling, resulting in a total sampling time of 100 ns for each FEP run in each direction (forward or backward). This process was repeated for both forward and backward directions for each of the two cation-bound states, totaling 400 ns for each mutation free energy calculation. The initial structures for all FEP simulations were equilibrated for at least 10 ns in the corresponding cation-binding states (Table 1), following the original 200 ns MD equilibration described above. The total FEP simulation time was ∼2 μs.

### PROPKA calculations of pKa’s of D55 and D59 residues

The absolute pKa’s of D55 and D59 in the WT mutants were estimated using the PROPKA program(46, 47). For each system, 2000 snapshots were selected from the three 200 ns MD production trajectories with a 0.1 ns interval. For each snapshot, the pKa’s of the D59 and D55 are calculated, and the results are averaged over all snapshots. Consistent with the simulation procedure used in ref. (14), for the estimation of D55’s pKa, the D59 residue is deprotonated and D55 is protonated throughout the MD simulation. For the estimation of D59’s pKa, the D59 residue is deprotonated and D55 is protonated in the simulation. No Na+ ion was present in the cation-binding site in all simulations.

### Statistics and reproducibility

All experiments were performed 2-4 times. The average values were presented in the table with standard errors. An unpaired t-test was used for statistical analysis.

## Abbreviations

MFS: major facilitator superfamily
ITC: isothermal titration calorimetry
FRET: fluorescence resonance energy transfer
FEP: free-energy perturbation
D^2^G, 2′: (*N*-dansyl)aminoalkyl-1-thio-β-D-galactopyranoside
α-NPG: 4-nitrophenyl-α-D-galactopyranoside
RSO membrane: right-side-out membrane
ΔΔ*G*_cation-binding_: free energy change for a cation binding upon a mutation
Δ*G*_mutation, cation-unbound state_: reflects the free energy change introduced by a mutation when MelB_St_ is at a cation-free form
Δ*G*_mutation, cation-bound state_: reflects the free energy change introduced by a mutation when MelB_St_ is at a cation-bound form

## Acknowledgments

We thank Gerard Leblanc for a MelB-expressing vector, the DW2 strain, and the dansyl galactoside. The x-ray diffraction datasets for the apo D59C and DDMB-bound D55C structures were collected at ALS BL 8.2.2 and 5.0.1, respectively. This work was supported by the National Institutes of Health Grant R01 GM122759 to L.G. and R35 GM150780 and Welch Foundation to R.L.

## Data availability statement

The x-ray diffraction datasets and models have been deposited to wwPDB under the accession code 8FQ9 for D55C MelB_St_ with DDMB and 8FRH for D59C MelB_St_ at an apo state.

