## Supplemental materials for "Distinct roles of the major binding residues in the cation-binding pocket of MelB"

**Supplementary Information Table S1**

Table S1. Setups of all FEP simulations involved in the relative binding affinity calculations. The initial structure column indicates the MD equilibrated variants of MelB_St_ (WT, D59C or D55C) as the starting point for the FEP calculation. See Table 2 for the definitions of free energy changes.

| Property evaluated | Mutation | Initial structure/  Simulation ID | Na^+^ bound? | D59 protonated? | D55 protonated? |
| --- | --- | --- | --- | --- | --- |
| ΔΔ*G*_Na+ binding_ | WT$\leftrightarrow$D55C | WT/1 | Yes | No | No |
|  |  | WT/2 | No | No | No |
|  | WT$\leftrightarrow$D59C | WT/1 | Yes | No | No |
|  |  | WT/2 | No | No | No |
|  | WT$\leftrightarrow$T121A | WT/1 | Yes | No | No |
|  |  | WT/2 | No | No | No |
| ΔΔ*G*_proton binding at Asp59_ | WT$\leftrightarrow$ D55C | D55C/1 | No | Yes | No |
|  |  | D55C/2 | No | No | No |
| ΔΔ*G*_proton binding at Asp55_ | WT$\leftrightarrow$ D59C | D59C/1 | No | No | Yes |
|  |  | D59C/2 | No | No | No |
